# Sensitization to Ionizing Radiation by MEK inhibition is Dependent on SNAI2 in Fusion-negative Rhabdomyosarcoma

**DOI:** 10.1101/2022.05.02.490343

**Authors:** Nicole R. Hensch, Kathryn Bondra, Long Wang, Prethish Sreenivas, Xiang R. Zhao, Paulomi Modi, Angelina V. Vaseva, Peter J. Houghton, Myron S. Ignatius

## Abstract

In fusion-negative rhabdomyosarcoma (FN-RMS), a pediatric malignancy with skeletal muscle characteristics, > 90% of high-risk patients have mutations that activate the RAS/MEK signaling pathway. We recently discovered that SNAI2, in addition to blocking myogenic differentiation downstream of MEK signaling in FN-RMS, represses pro-apoptotic *BIM* expression to protect RMS tumors from ionizing radiation (IR). As clinically relevant concentrations of the MEK inhibitor trametinib elicit poor responses in preclinical xenograft models, we investigated the utility of low-dose trametinib in combination with IR for the treatment of RAS-mutant FN-RMS. We hypothesized that trametinib would sensitize FN-RMS to IR through its downregulation of SNAI2 expression. While we observed little to no difference in myogenic differentiation or cell survival with trametinib treatment alone, robust differentiation and reduced survival were observed after IR. Additionally, IR-induced apoptosis was significantly increased in FN-RMS cells treated concurrently with trametinib, as was increased BIM expression. SNAI2’s role in these processes was established using overexpression rescue experiments, where overexpression of SNAI2 prevented IR-induced myogenic differentiation and apoptosis. Moreover, combining MEK inhibitor with IR resulted in complete tumor regression and a 2–4-week delay in event free survival (EFS) in preclinical xenograft and PDX models. Our findings demonstrate that the combination of MEK inhibition and IR results in robust differentiation and apoptosis, due to the reduction of SNAI2, which leads to extended EFS in FN-RMS. SNAI2 thus is a biomarker of IR insensitivity and target for future therapies to sensitize aggressive sarcomas to IR.

## Introduction

Rhabdomyosarcoma (RMS) is a pediatric malignancy resembling skeletal muscle. There are two major clinical sub-types: fusion-positive (FP) RMS, driven by either PAX3-FOXO or PAX7-FOXO fusions, and fusion-negative (FN) RMS, where genomic sequencing studies confirmed > 90% high risk patients have RAS-activating mutations present^1–4^. While the outcome for the FN-RMS subtype is often considered more favorable than FP-RMS, tumors with aberrant MEK signaling caused by *RAS* mutations have been shown to be more aggressive and difficult to treat with mainstay therapies^1–3^. Moreover, the 5-year survival in patients with relapse or metastasis is less than 30%^5–8^. For the pediatric patients diagnosed with RMS, the current standard of care includes surgery, chemotherapy, and ionizing radiation (IR)^9–11^. Currently, there are no FDA-approved targeted therapies for the treatment of RMS^12^. Because aggressive tumors continue to be poor responders to conventional treatments, it is imperative to discover novel therapeutic targets for FN-RMS to ensure better recurrence-free survival.

While there are no approved targeted therapies for RAS-mutant FN-RMS, recent studies suggest that inhibition of the RAS/MEK signaling cascade might be beneficial in the clinic^13, 14^. Trametinib and other MEK inhibitors have been identified as potential agents to target MEK1/2 in RAS-mutant RMS^12^. Yohe *et al.* demonstrated that treatment of FN-RMS cells and xenografts with high dose trametinib can result in RAS pathway inhibition and a delay in tumor growth^14^. However, while mice can tolerate higher doses of trametinib (3 mg/kg), we have determined a lower dose of trametinib leads to clinically relevant exposures, and at this dose the effect of trametinib inhibition of RAS/MEK signaling is less effective^13^. Recently, we showed that vertical inhibition of RAS/MEK/ERK signaling by combining trametinib with RAF inhibition is able to effectively inhibit MEK and ERK signaling and achieve complete tumor responses and enhanced delay in event free survival in preclinical models compared to single agent trametinib^13^. Nevertheless, while promising, whether combination treatments are tolerated and tumor responses are maintained after vertical inhibition of the RAS/MEK/ERK pathway in patients remains to be determined. Another promising avenue would be to utilize MEK inhibition (MEKi) with a standard of care therapy. There are several active clinical trials investigating the utility of MEKi in combination with IR in other cancers, including non-small cell lung cancer, melanoma, and brain metastases^15–17^. However, the exact mechanisms and effectors by which hyperactive RAS/MEK signaling protects tumors from standard of care treatments have not been fully elucidated.

Recently, we discovered that transcriptional repressor SNAI2, which is highly expressed in all RMS tumors, plays a pivotal role in protecting cells from IR-induced apoptosis by directly repressing the expression of pro-apoptotic regulator *BIM* (*BCL2L11)*^18^. Independently, in RAS-mutant FN-RMS cells and xenografts, SNAI2 is also a potent repressor of myogenic differentiation acting through multiple mechanisms^19^. We also discovered that in FN-RMS cells, RAS-MEK signaling was required to maintain SNAI2 protein expression^19^. Interestingly, there was a high overlap in the genes modulated by SNAI2 and RAS-MEK signaling, suggesting that SNAI2 may be an important effector of RAS/MEK signaling effects on myogenic differentiation^19^. Additionally, BIM is known to be post-translationally regulated by MEK/ERK signaling, suggesting a convergence of effects on preventing apoptosis^20^. In this study, we sought to address if combining low-dose trametinib with IR would be effective in treating RAS-mutant FN-RMS tumors (Supplemental Table 1). Additionally, we investigated whether SNAI2 is the critical downstream effector of RAS/MEK signaling that prevents myogenic differentiation and protects FN-RMS tumors from radiation. Our findings indicate that the reduction of SNAI2 is responsible for the sensitization of FN-RMS when treated with IR and trametinib. FN-RMS pre-treated with trametinib behaved similarly to that of SNAI2 knockdown RMS cells previously characterized, in that myogenic differentiation was increased and overall survival post IR was reduced. Additionally, a larger population of cells were undergoing apoptosis after combined trametinib and IR treatment when compared to cells treated with either single-agent therapy. In overexpression assays, increasing SNAI2 in FN-RMS cells is sufficient to reduce IR-induced apoptosis and to block myogenic differentiation induced by the combination of trametinib and IR treatment. Finally, in RAS-mutant xenograft/PDX-bearing mice, trametinib treatment led to reductions of SNAI2, and when combined with IR treatment resulted in complete responses and increased event free survival in each of the three independent FN-RMS xenograft models. Our analysis suggests that the Ras/MEK/ERK signaling pathway, through the maintenance of SNAI2 expression, protects FN-RMS against apoptosis and differentiation post IR and that reducing SNAI2 by targeting the MEK/SNAI2 axis could be a viable solution to increase tumor-killing responses to IR therapies in FN-RMS patients.

## Materials and Methods

### Human rhabdomyosarcoma (RMS) cell lines

The human RMS cell lines RD were a gift from Dr. Corinne Linardic, Duke University, Durham, NC. The SMS-CTR cell line was a gift from Dr. Angelina Vaseva, GCCRI, San Antonio, TX. The Rh36 cell line was a gift from Dr. Peter J. Houghton, GCCRI, San Antonio, TX. SMS-CTR and Rh36 cells were maintained in RPMI supplemented with 10% fetal bovine serum (VWR) at 37° C with 5% CO2. RD cells were maintained in DMEM supplemented with 10% FBS at 37° C with 5% CO2. Cell lines utilized were between passage 10 and 25. Cell lines were authenticated by genotyping and STR analyses. All RMS cell lines were tested and confirmed to be negative for mycoplasma.

### Western blot analysis

Total cell lysates from human RMS cells and xenografts were obtained following lysis in SDS lysis buffer. Western blot analysis was performed similar to Ignatius et. al., 2017^21^. Membranes were developed using an ECL reagent (Western Lightning Plus ECL, PerkinElmer; or sensitive SuperSignal West Femto Maximum Sensitivity Substrate, Thermo Scientific). Membranes were stripped, rinsed, and re-probed with the respective internal control antibodies. List of primary and secondary antibodies is included in supplementary data (Supplemental Table 2).

### Immunofluorescence Staining

Immunofluorescence staining was performed as described in Wang *et al*., 2021^18^. Cells were pretreated with DMSO or trametinib (10 nM for 72h or 20 nM for 24h) prior to being plated at 4,000 cells/well (no IR) and 10,000 cells/well (receiving IR), grown in 10% FBS DMEM or RPMI growth media, fixed at 72 hours post IR (hpIR) (0 or 15 Gy) in 4% paraformaldehyde/PBS, permeabilized in 0.5% Triton X-100/PBS, and incubated with rabbit anti-MEF2C (CST; Catalog No. 5030) and anti-myosin heavy chain (DSHB) in 1% BSA/PBS. Secondary antibody detection was performed with Alexa Flour 488 goat anti-mouse and Alexa Fluor 594 goat anti-rabbit (Invitrogen). Cells were counterstained with DAPI (1:10,000) and imaged. Images were processed in ImageJ and Adobe Photoshop. Significance was calculated by a two-way ANOVA with a posthoc Sidak’s multiple comparison test.

### Colony formation and cell confluency assays

For colony formation assays, RD cells (parental, pBabe, and SNAI2-Flag) were pretreated with either DMSO or trametinib (10 nM for 72h or 20 nM for 24h) prior to being seeded in 12-well plates (∼1,200–20,000 cells/well for cells receiving radiation and 300 cells/well for cell receiving no IR). After 24h, cells were subjected to varying degrees of radiation (0–6 Gy). Incubation time for colony formation assays between cell lines varied from 1 to 6 weeks. When colonies were sufficiently large, media was gently removed from each plate by aspiration, and colonies were fixed with 50% methanol for 10–15 minutes at room temperature (RT). Colonies were then stained with 3% (w/v) crystal violet in 25% methanol for 10–15 minutes RT, and excess crystal violet was washed with dH2O with plates being allowed to dry. Colony formation was analyzed using ImageJ (Fiji) and significance for each radiation dose was calculated by Student’s t-test. RD and SMS-CTR parental cells were pretreated with either DMSO or trametinib (10 nM for 72h or 20 nM for 24h) prior to being seeded into 48-well plates at 20-50% confluency and stored at 37° C in the Incucyte ZOOM (Essen Bioscience). After 24h, cells were subjected to varying degrees of IR (0 Gy or 15 Gy), began respective DMSO or trametinib (10 nM) treatment, and placed back in the Incucyte. Total confluency over time was monitored every 4–6 hours over a period of 5 days. Significance was calculated by one-way ANOVA with Dunnett’s multiple comparisons tests.

### Flow Cytometry

RD and SMS-CTR cells were seeded in 100 mm plates and treated with DMSO or trametinib (10 nM for 72h or 20 nM for 24h). After sufficient DMSO or trametinib treatment, cells were irradiated (PXi Precision X-Ray X-RAD 320) and continued DMSO or trametinib (10 nM) treatment for 72h until cells were collected. A set of DMSO and trametinib-treated cells that did not receive IR were collected as well. Cells were centrifuged and resuspended in annexin-binding buffer. After determining cell density and diluting to 1 × 10^6^ cells/mL with annexin-binding buffer, annexin V conjugate, and propidium iodide were added to sample aliquots and left to incubate at room temperature in the dark for 15 minutes. After incubation, aliquots were mixed gently while adding annexin-binding buffer on ice and analyzed by flow cytometry (LSRFortessa X-20; BD Biosciences). Cell cycle was assessed using the same cells and conditions described above with Click-iT EdU Alexa Fluor 647 Flow Cytometry Assay (ThermoFisher) according to the provided protocol. Significance was determined using a Two-Proportion Z-test. All assays were performed in triplicates.

### ChIP-seq

ChIP-seq tracks of antibodies targeting SNAI2 (CST, Catalogue # 9585), and H3K27ac (Active Motif, cat. #39133) were used for the current analysis (GEO datasets GSE137168, GSE85171)^14, 19^. ChIP-seq fastq files were mapped to human reference genome (hg19) using Bowtie2. High-confidence ChIP-seq peaks were called by MACS2 using default parameters and peaks were visualized using IGV viewer or epigenome browser.

### Retroviral Expression assays

Control and *SNAI2*-specific Flag tag were delivered via the pBabe-background vector and packaged using 293T cells. pPGS-hSLUG.fl.flag was a gift from Eric Fearon (Addgene plasmid # 25696 ; http://n2t.net/addgene:25696; RRID:Addgene_25696). Retroviral particles were made in Plat-A packaging cells using TranstIT-LT1 (Mirus). RMS cells were infected with viral particles for 24h at 37° C using 8 mg/mL of polybrene (EMD Millipore). Cells were selected for using 1 mg/ml G418 (Sigma, Cat. G8168) for 7 days.

### Animal Studies

Animal studies were approved by the University of Texas – Health San Antonio Committee on Research Animal Care under protocol #20150015AR (Peter Houghton). C.B-Igh-1b/IcrTac-Prkdcscid (SCID) female mice, aged 6–8 weeks (Envigo, Indianapolis, IN), were used for *in vivo* xenograft experiments.

### Mouse xenograft and *in vivo* IR experiments

SJRHB013_X and SJRHB000026_X1 patient-derived xenograft (PDX) models were obtained from the Childhood Solid Tumor Network^22^. Xenografts were transplanted into the left hind left leg of the mice and left to grow to a sufficient size to collect and engraft equal size tumor pieces into approximately 45 mice each. RD cells were collected, counted, and analyzed by flow cytometry to determine viability using DAPI. Equal numbers of viable cells were then embedded into Matrigel at a final concentration of 1×10^6^ (RD) per 100 μl and injected subcutaneously into anesthetized mice. Tumors were allowed to grow to a sufficient size to collect and engraft equal size tumor pieces into approximately 45 mice. Once tumors reach approximately 0.2 cm^3^ in size, tumor growth was monitored and measured weekly using a caliper scale to measure the greatest diameter and length, which were then used to calculate tumor volume. While a subset of tumors was monitored without any treatment, another subset was treated with continuous 28-day treatment with trametinib (1 mg/kg) via oral gavage. A third treatment arm was subjected to IR therapy for 1–3 weeks (2 Gy/day; 5 days a week), receiving a total of 10-Gy (SJRHB013_X), 20-Gy (SJRHB000026_X1), or 30-Gy IR (RD) (PXi Precision X-Ray X-RAD 320). IR dosages were determined after an IR sensitivity study was conducted, where 3–6 xenograft-burdened mice were subjected to either 10, 20, or 30 Gy of radiation and measured for relative tumor volume (RTV) over time. The last treatment group included both 28-day trametinib treatment with IR therapy, where trametinib treatment started 3–5 days before IR therapy. Tumor volume was monitored throughout the treatment and during the weeks following treatment. Comparisons between groups was performed using a one-way ANOVA with a posthoc Tukey test. RTV was assessed in the RD, SJRHB013_X, and SJRHB000026_X1 tumors, and tumor responses were analyzed using the guidelines outlined by Houghton *et al*.^23^. Progressive disease (PD): RTV > 0.50 during the study period and RTV > 1.25 at end of study; Stable disease (SD): RTV > 0.50 during the study period and RTV ≤ 1.25 at end of study; Partial response (PR): 0 < RTV < 0.50 for at least one time point; Complete response (CR): RTV = 0 for at least one time point. Average event-free survival (EFS) was calculated as the mean number of weeks it took for tumors to reach 4 times their initial tumor volume.

## Results

### Low-dose MEK inhibitor reduces SNAI2 expression and in combination with IR induces robust myogenic differentiation and decreased cell survival in FN-RMS

FN-RMS are often driven by aberrant RAS/MEK signaling, and mutations that activate the RAS/MEK signaling pathway are frequently observed in this sub-type. Previously, we demonstrated that trametinib, a potent MEK inhibitor, reduced SNAI2 in FN-RMS cell lines in addition to inhibiting MEK/ERK signaling^19^. To confirm this assessment, we treated FN-RMS parental cells with DMSO and low-dose trametinib (10 nM) and assessed protein expression in multiple FN-RMS cell lines. As expected, RD, SMS-CTR, and Rh36 cells displayed reduced SNAI2 expression following trametinib treatment compared to control treated cells. Additionally, we observed that trametinib-treated cells also had increased BIM expression in comparison to control cells treated with DMSO (**Figures 1A–C**). In contrast, trametinib treatment increases SNAI1 protein expression, consistent with increased SNAI1 resulting from loss of SNAI2. Thus, the RAS-MEK-ERK pathway is required for SNAI2 expression, while at the same time represses pro-apoptotic BIM expression in FN-RMS.

**Figure 1.**
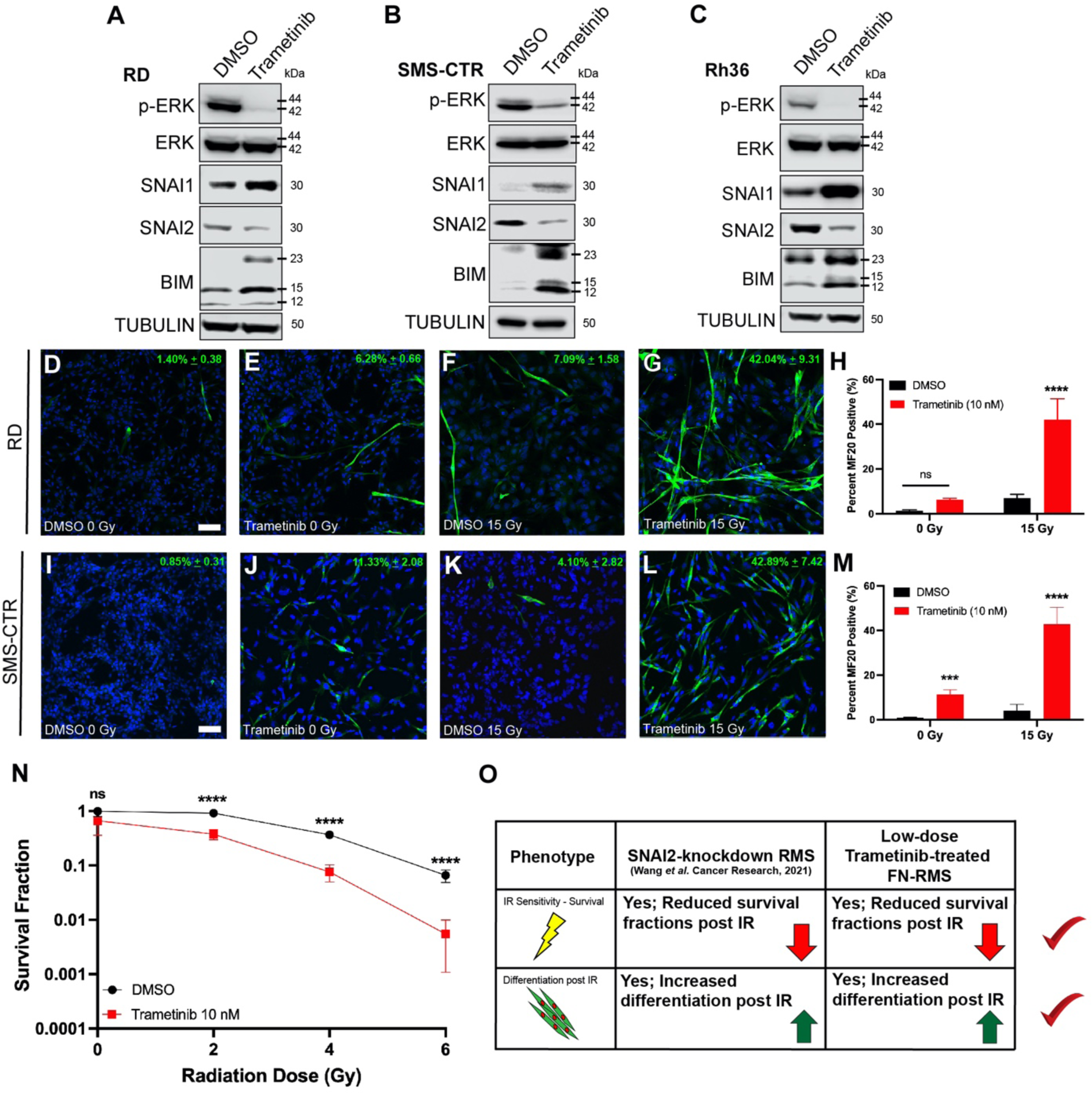
Low-dose MEK inhibitor reduces SNAI2 expression and in combination with IR induces robust myogenic differentiation and loss of cell survival in FN-RMS. A, B, C. Protein expression in DMSO- and trametinib-treated FN-RMS cell lines (RD, SMS-CTR, and Rh36). D-G. Representative confocal microscopy images of RD cells treated with DMSO or trametinib, in non-IR (D, E) and IR (F, G) conditions, immunostained with myogenic differentiated myosin (MF20 antibody). Scale bar = 100 μm. H. Quantification of average MF20-positive cells in either non-IR or IR conditions in DMSO- and trametinib- treated RD cells. Error bars represent ±1 SD. ns = not significant, \*\*\*\**p*<0.0001 by a two-way ANOVA with a posthoc Sidak’s multiple comparison test. I–L. Representative confocal microscopy images of SMS-CTR cells treated with DMSO or trametinib, in non-IR (H, I) and IR (J, K) conditions, immunostained with myogenic differentiated myosin (MF20 antibody). Scale bar = 100 μm. M. Quantification of average MF20-positive cells in either non-IR or IR conditions in DMSO- and trametinib-treated SMS-CTR cells. Error bars represent ±1 SD. ns = not, \*\*\**p* < 0.001, \*\*\*\**p* < 0.0001 by a two-way ANOVA with a posthoc Sidak’s multiple comparison test. N. Survival fractions of DMSO- and trametinib-treated RD cells were assessed at increasing IR dose exposures. Statistical differences were observed at 2, 4, and 6 Gy. Error bars represent ±1 SD. ns = not significant, \*\*\*\**p* < 0.0001 by a Student’s t-test. O. Overview of similar phenotypic characteristics of SNAI2-knockdown RMS cells and low-dose trametinib- treated FN-RMS cells.

SNAI2 is a potent inhibitor of myogenic differentiation and combining SNAI2 knockdown with ionizing radiation (IR) results in robust myogenic differentiation in addition to loss of colony forming ability and increased apoptosis^18^. Therefore, we tested if combining treatment of low-dose trametinib and IR in RD and SMS-CTR cells would lead to similar increased differentiation as in SNAI2-shRNA knockdown FN-RMS cells. We performed differentiation assays assessing myogenic differentiation after trametinib treatment with or without IR exposure in growth media rather than in low serum media that is often used to initiate myogenic differentiation ^21, 24–26^. FN-RMS cells were treated with DMSO or trametinib, and a subset of cells were also subjected to IR. RD cells that received trametinib treatment but were not exposed to IR had slightly higher levels of differentiation compared to the DMSO controls, although not statistically significant (*p* = 0.1703, **Figures 1D, E**). However, RD cells that received IR after trametinib pre-treatment underwent robust myogenic differentiation compared to IR-exposed, DMSO-treated cells (*p* < 0.0001, **Figures 1F–H**). SMS-CTR cells were even more sensitive to trametinib treatments, and while both non-IR and IR-exposed SMS-CTR cells treated with trametinib showed significantly higher myogenic differentiation compared to their respective DMSO controls, the trametinib + IR combination resulted in more robust differentiation compared to trametinib alone (42.89% ± 7.42 vs.11.33% ± 2.08, *p* = 0.0005 (0 Gy), *p* < 0.0001 (15 Gy), **Figures 1I–M**).

To evaluate the effect of combining trametinib with IR on cell survival, we performed colony forming assays in DMSO- and trametinib-treated RD cells. Cells were treated with DMSO or trametinib (10 nM) prior to plating and radiation exposure. In the absence of IR, there was no statistical difference in survival in DMSO- and trametinib-treated cells. However, upon IR exposure, we noted a significant difference at all IR doses with cells having received trametinib treatment showing decreased survival fractions compared to DMSO controls (*p* < 0.0001 for 2–6 Gy, **Figure 1N**, Supplemental Figure 1A). SMS-CTR cells were also studied; however, their sensitivity to trametinib led to poor plate attachment and colony forming units could not be accurately quantified. Additionally, we also assessed RD and SMS-CTR cell confluency to determine the effects of combining trametinib and IR. Indeed, while RD was overall less sensitive to either trametinib or IR alone, a significant drop in confluency was observed after combination therapy (Supplemental Figure 1B). As expected, SMS-CTR cells exhibited higher sensitivities to both trametinib and IR; and the combination of the two significantly reduces confluency compared to single agent treatments (Supplemental Figure 1C). Together, our analyses show that MEK signaling is required for SNAI2 expression and that combining a low dose of MEK inhibitor, trametinib, with IR, reduces SNAI2 and sensitizes cells to IR causing increased myogenic differentiation and reducing cell survival, that recapitulate the phenotypes found in SNAI2 shRNA knockdown RMS cells (**Figure 1O**).

### Combining low-dose MEK inhibitor with IR results in increased apoptosis and G1 cell cycle arrest in FN- RMS

SNAI2 knockdown RMS cells are known to undergo IR-induced apoptosis. Therefore, to determine the effect of combining low dose trametinib and IR treatment on apoptosis, we utilized flow cytometric analysis with Annexin V and propidium iodide staining. As expected, we observed that there was little to no apoptosis in DMSO-treated cells without IR exposure and increased apoptosis after treatment with trametinib in RD cells (**Figures 2A, B**). However, after IR exposure (15 Gy), trametinib-treated cells underwent significantly increased apoptosis compared to their DMSO-treated counterparts (*p* < 0.0001, **Figures 2C, D**). SMS-CTR cells are more sensitive to continuous trametinib treatment that results in approximately 30% of cells undergoing early to late apoptosis compared to the ∼6% of cells in DMSO treatment alone (**Figures 2E, F**). SMS-CTR cells that received 10 Gy of IR with continuous DMSO demonstrated a ∼20% population of cells undergoing apoptosis; however, continuously trametinib-treated cells that received the same IR dose led to an increased fraction of cells undergoing apoptosis (∼47%), suggesting there is a significant additive effect of co-treating FN-RMS cells with trametinib and IR (*p* < 0.0001, **Figures 2G, H**). These findings indicate that trametinib treatment sensitizes FN-RMS to undergo radiation-induced apoptosis.

**Figure 2.**
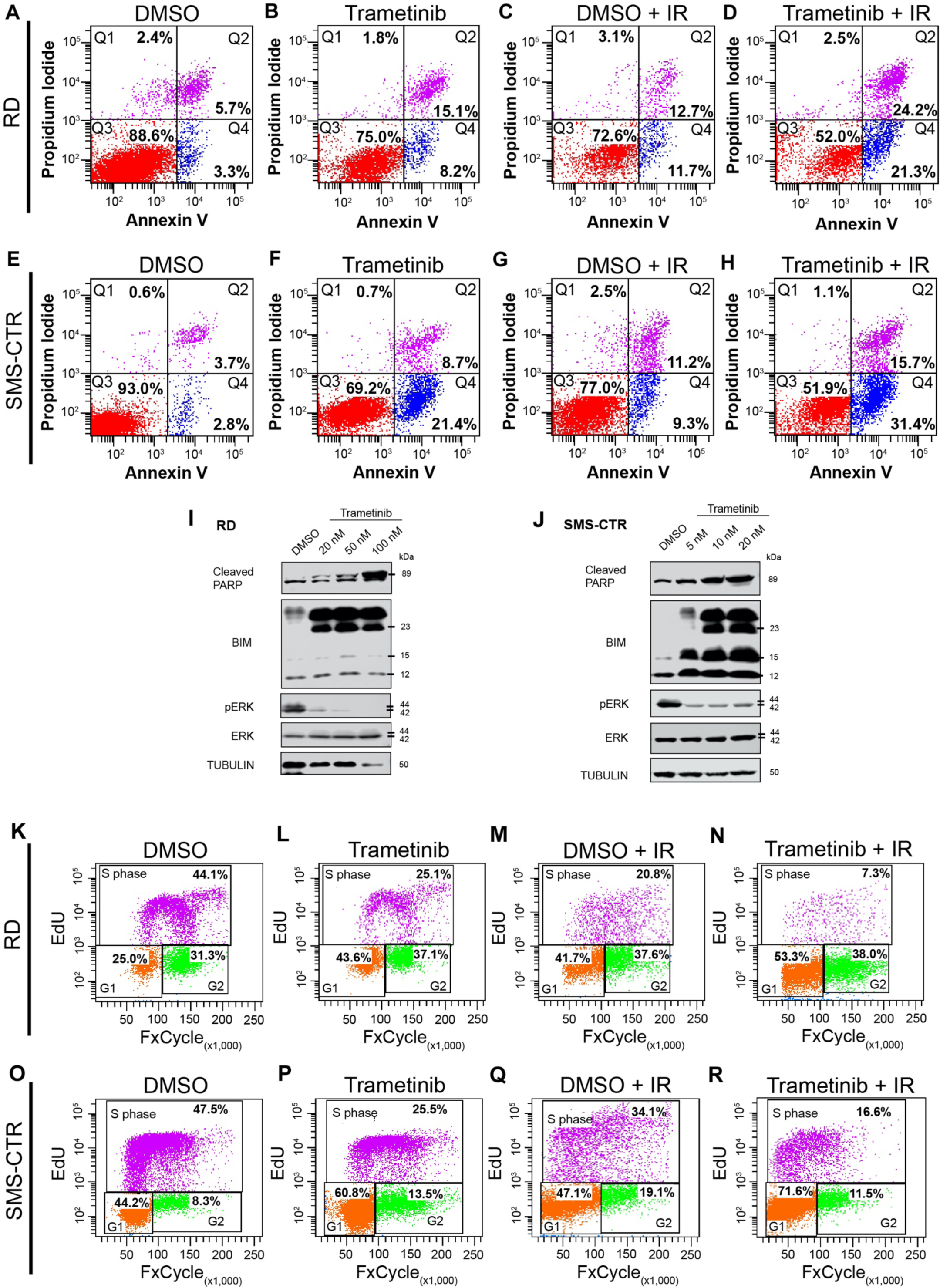
Combining low-dose MEK inhibition with IR results in increased apoptosis and a G1 cell cycle arrest in FN-RMS. A-D. Flow cytometry plots showing propidium iodide vs. annexin V staining of DMSO- and trametinib-treated RD cells after non-IR (A, B) and IR (C, D) conditions. Q4 represents cells undergoing early apoptosis, whereas Q2 represents cells undergoing late apoptosis. Q3 represents live cells not undergoing apoptosis. E-H. Flow cytometry plots showing propidium iodide vs. annexin V staining of DMSO- and trametinib-treated SMS-CTR cells after non-IR (E, F) and IR (G, H) conditions. I, J. Cleaved PARP, BIM, pERK, and total ERK protein expression immunoblots in DMSO- and trametinib-treated RD (I) and SMS-CTR (J) cells 96hpIR. Increased trametinib doses were assessed in both cell lines. K-N. Flow cytometry plots of EdU vs. DAPI staining in DMSO- and trametinib-treated RD cells in non-IR (K, L) and IR (M, N) conditions. O-R. Flow cytometry plots of EdU vs. DAPI staining in DMSO- and trametinib-treated SMS-CTR cells in non-IR (O, P) and IR (Q, R) conditions.

To investigate the mechanism by which combination trametinib and IR treatment induces apoptosis, we assessed protein expression for the terminal apoptosis marker, cleaved PARP, and pro-apoptotic BIM after the various treatments, as we have previously shown that SNAI2-knockdown RMS cells undergo apoptosis via a BIM-mediated mechanism after IR exposure^18^. Trametinib treatment resulted in reduced SNAI2, a slight increase in cleaved PARP, and increased expression of BIM prior to radiation exposure, suggesting these cells are primed for radiation-induced apoptosis as previously shown^18^ (Supplemental Figure 2A, B). After combining IR with continuous trametinib treatment RD and SMS-CTR cells both demonstrated high levels of cleaved PARP and BIM compared to DMSO-treated cells post IR, with cleaved PARP increasing as trametinib concentrations increased (**Figures 2I, J**). We previously showed that SNAI2 can directly repress *BIM* expression in RMS by engaging two enhancer elements^18^. Using published data for SNAI2 binding post trametinib treatment in FN-RMS, we show that SNAI2 binding at the more distal 3’ enhancer is lost after 48-hour trametinib treatment (Supplemental Figure 2C).

We next assessed the effect of combining trametinib and IR on the cell cycle. Compared to control RD cells, a significant number of trametinib-treated cells accumulated in the G1 phase (**Figures 2K, L**). While a similar G1 increase was noted in the DMSO-treated cells after IR exposure, this was exacerbated in the cells receiving combination treatment, with a significant decrease in actively dividing cells also apparent (*p* < 0.0001, **Figures 2M, N**). Similar observations were also seen in SMS-CTR cells under in the same treatment conditions (*p* < 0.0001, **Figures 2O–R**), indicating that trametinib either alone or in combination with IR results in a G1 cell cycle block in FN-RMS cells. These data do not correlate with SNAI2-knockdown, which results in a G2/M accumulation, suggesting that the effects of trametinib on the cell cycle may be independent of SNAI2 downregulation.

### SNAI2 downstream of MEK signaling protects FN-RMS from IR-induced myogenic differentiation and apoptosis

Our experiments indicate that a MEK/SNAI2 signaling axis protects RAS-mutant FN-RMS cells from IR by blocking myogenic differentiation and preventing apoptosis. We therefore performed epistasis experiments to determine if re-expressing SNAI2 in MEK inhibitor-treated cells can protect cells from IR by preventing myogenic differentiation and apoptosis. We first generated retroviral-induced stable SNAI2 overexpressing (SNAI2-Flag) RD and SMS-CTR cells (**Figure 3A**, Supplemental Figure 3A). We next tested the effect of overexpressing SNAI2 on myogenic differentiation post combining low dose trametinib with IR. Whereas the RD control cells that were treated with trametinib displayed a statistically significant increase in terminal differentiation, the RD SNAI2-Flag trametinib-treated cells had noticeably less terminal myogenic differentiation than their control counterparts (*p* < 0.0001, **Figure 3B**). Similarly, SMS-CTR pBabe control and SNAI2-Flag cells that were treated with trametinib also resulted in increased differentiation post-IR, similar to the studies done in parental cells, the SMS-CTR SNAI2-Flag cells had significantly less cells positive for myogenic differentiation markers when compared to SMS-CTR pBabe controls (*p* < 0.0001, Supplemental Figure 3B).

**Figure 3.**
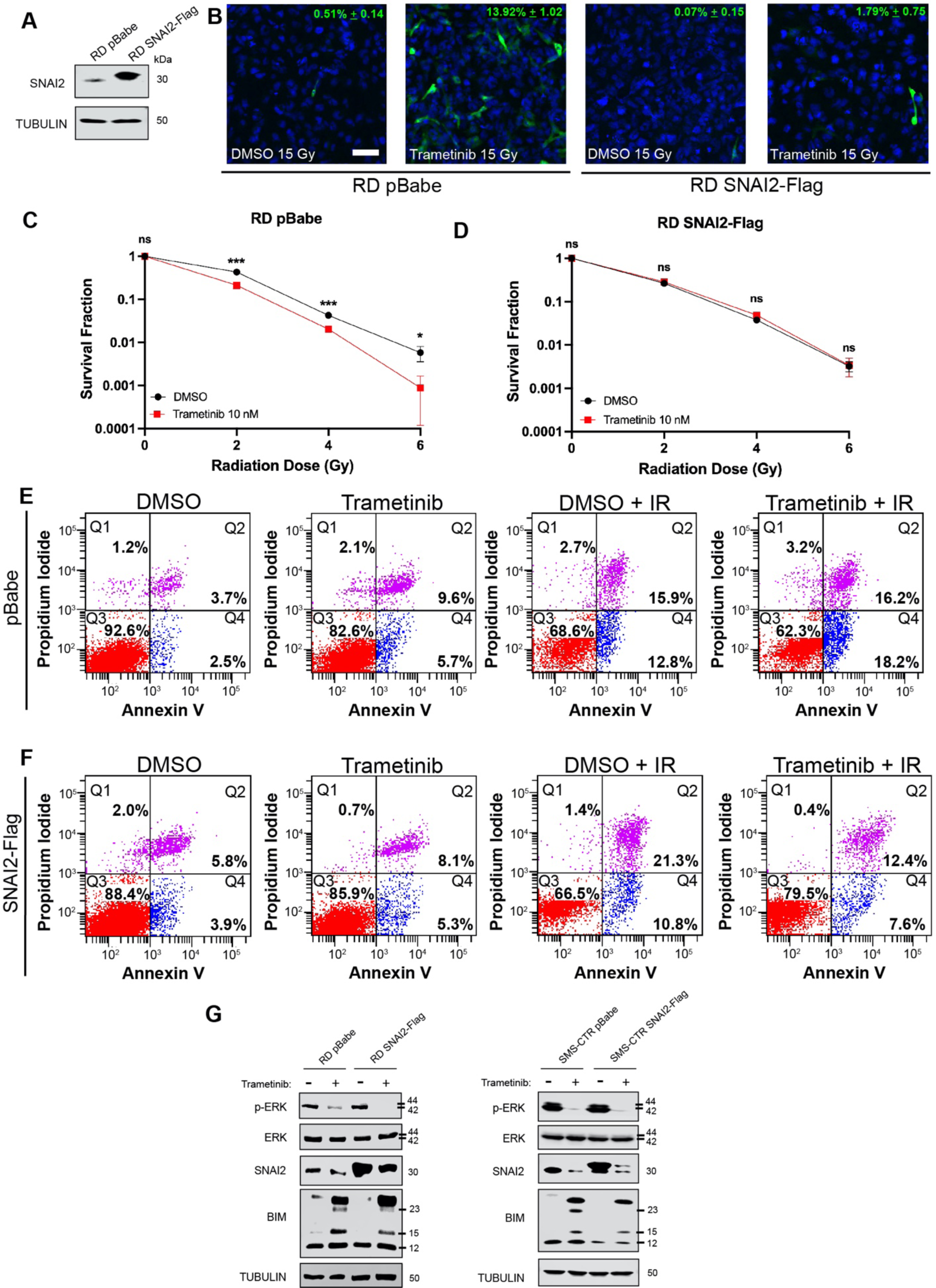
SNAI2 downstream of MEK signaling protects FN-RMS from IR-induced myogenic differentiation and apoptosis. A. SNAI2 protein expression in RD pBabe and SNAI2-Flag overexpressing cell lines. B. Representative confocal microscopy images of RD pBabe and SNAI2-Flag cells treated with DMSO or trametinib after IR exposure, immunostained with myogenic differentiated myosin MF20 antibody. Scale bar = 100 μm. C, D. Survival fractions of DMSO- and trametinib-treated RD pBabe and SNAI2-Flag cells were assessed at increasing IR dose exposures. Statistical differences were observed at 2, 4, and 6 Gy in RD pBabe, but NOT RD SNAI2-Flag. Error bars represent ±1 SD. ns = not significant, \**p* < 0.05, \*\*\**p* < 0.001 by a Student’s t-test. E. Flow cytometry plots showing propidium iodide vs. annexin V staining of DMSO- and trametinib-treated RD pBabe cells after non-IR and IR conditions. F. Flow cytometry plots showing propidium iodide vs. annexin V staining of DMSO- and trametinib-treated RD SNAI2-Flag cells after non-IR and IR conditions. G. Western blots showing pERK, total ERK, and BIM protein expression of RD pBabe, RD SNAI2-Flag, SMS- CTR pBabe, and SMS-CTR SNAI2-Flag cells after DMSO and 10 nM trametinib treatment.

We then evaluated the effect of SNAI2 overexpression on clonal growth post combination with trametinib and IR. While we observed the expected separation of trametinib + IR-treated control cells in terms of survival fraction, this decrease in survival was lost in SNAI2-Flag overexpressing cells, indicating that loss of SNAI2 downstream of MEK is responsible for the sensitization to IR in parental/control cells (**Figures 3C, D**).

Next, we examined whether the abundance of SNAI2 in trametinib and IR-treated SNAI2 overexpressing cells would prevent the significantly increased apoptosis seen in FN-RMS RD and SMS-CTR cells. As expected, the RD cells that received combinatorial treatment had increased levels of early and late apoptosis, as determined by Annexin V staining, compared to either single agent treatment (**Figure 3E**). However, while the trametinib and IR single agent exposure did not result in statistically different apoptosis levels, the RD SNAI2-Flag cells had significantly reduced apoptosis after combining trametinib and IR (Q2 and Q4 populations, *p* < 0.0001, **Figure 3F**). SMS-CTR cells remained sensitive to combinatorial low-dose trametinib + IR conditions, resulting in increased apoptosis (Q2 and Q4 populations, *p* < 0.0001, Supplemental Figure 3C). In contrast, no overall differences in apoptosis (early + late apoptosis) were observed in SMS-CTR SNAI2-Flag cells receiving low-dose trametinib treatment compared to DMSO-treated cells, regardless of IR treatment status (Supplemental Figure 3D). This reduction in apoptosis implies that the overexpression of SNAI2 prevents apoptosis post trametinib and IR treatment. Finally, we tested if the overexpression of SNAI2 prevents BIM expression in the trametinib treatment setting. We assessed in trametinib-treated cells if the overexpression of SNAI2 resulted in the loss of BIM expression and show that SNAI2 overexpression is able to represses BIM expression after trametinib treatment, while in control trametinib-treated cells, BIM is robustly induced (**Figure 3G**). Together, these experiments demonstrate that SNAI2 downstream of MEK signaling protects RAS mutant FN-RMS cells from IR by preventing myogenic differentiation and apoptosis.

### MEK inhibitor treatment results in loss of SNAI2 and increases sensitivity to IR, resulting in increased event-free survival in preclinical models

Our data thus far indicate that SNAI2 loss is likely a good biomarker and key determinant for response to the combination of MEK inhibition with IR. To assess the efficacy of combining MEK inhibition, via trametinib, with IR treatment in preclinical models *in vivo*, we tested four treatment arms: control (untreated), trametinib (1 mg/kg x 28 days), IR (10–30 Gy total; 2 Gy/day x 5 days a week until complete), and combination of trametinib and IR in RAS-mutant (mutation) RD xenografts and two RAS-mutant FN-RMS PDXs (**Figure 4A**). In control IR experiments, we initially determined the sensitivity of xenografts to 10, 20 and 30 Gy IR on loss of tumor volume and relapse (n = 3–5 mice/treatment) and found that RD xenografts are resistant to even 20 Gy of IR, SJRHB000026_X1 (RAS mutation) partially respond to 20 Gy, while SJRHB013_X (RAS mutation) was relatively sensitive to IR (Supplemental Figure 2A–C)^1^. Based on these findings, we selected IR increments of 10 Gy (SJRHB013_X – 10 Gy, SJRHB000026_X1 – 20 Gy, and RD – 30 Gy). After tumors reached approximately 0.2 cm^3^, mice were randomly separated into the four treatment groups. Tumors that received oral trametinib for ∼5 days were characterized by marked reduction of pERK and SNAI2, in addition to increased BIM protein expression indicating that our dosing schedule was effective at inhibiting RAS/MEK signaling (**Figures 4B–D**). Control RD xenografts rapidly grew to 4x their initial volume in 2 weeks (**Figure 4E**). Utilizing the clinically relevant trametinib low-dose treatment extended the average event free survival (EFS) to 4.7 weeks, while RD xenografts that received 30 Gy IR averaged an EFS of 13.4 weeks (**Figures 4F, G**). However, in combination, trametinib and IR treatment yielded an average EFS of 16.1 weeks, significantly delaying overall relapse of the tumor (*p* < 0.0055, **Figure 4H**).

**Figure 4.**
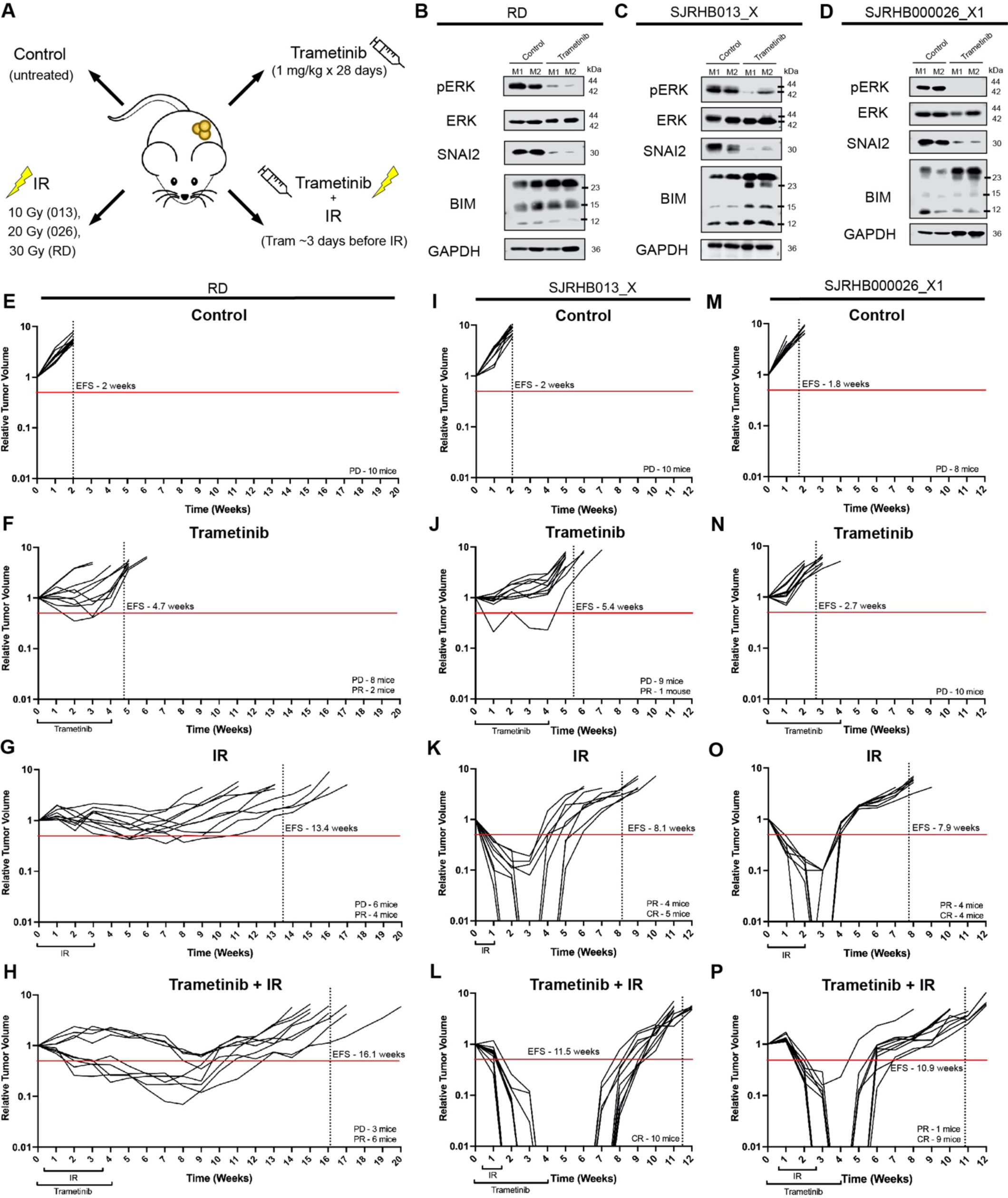
MEK inhibitor treatment results in loss of SNAI2 and increases sensitivity to IR, resulting in increased event-free survival in preclinical models. A. Illustration of *in vivo* experimental treatment set up. B–D. Western blots showing pERK, total ERK, SNAI2, and BIM protein expression in two representative control tumors and trametinib-treated RD (B), SJRHB013_X (C), and SJRHB000026_X1 (D) tumors. E–H. Relative tumor volume (RTV) of RD tumors receiving control (E), trametinib (F), IR (G), or combination trametinib + IR (H) treatment. Red line depicts 0.5 RTV; vertical dashed line represents the average EFS for each treatment arm. PD: progressive disease, PR: partial response. I–L. Relative tumor volume (RTV) of SJRHB013_X tumors receiving control (I), trametinib (J), IR (K), or combination trametinib + IR (L) treatment. Red line depicts 0.5 RTV; vertical dashed line represents the average EFS for each treatment arm. PD: progressive disease, PR: partial response. CR: complete response. M–P. Relative tumor volume (RTV) of SJRHB000026_X1 tumors receiving control (M), trametinib (N), IR (O), or combination trametinib + IR (P) treatment. Red line depicts 0.5 RTV; vertical dashed line represents the average EFS for each treatment arm. PD: progressive disease, PR: partial response. CR: complete response.

While combination treatment RD xenografts resulted in 6 out of 9 mice displaying partial responses (PR), more dramatic results were observed in the other two FN-RMS PDXs we assessed. SJRHB013_X control PDXs, when left untreated, reached 4 times their initial tumor volume after 2 weeks (**Figure 4I**). The EFS of SJRHB013_X tumor-burdened mice treated with trametinib was delayed by over 3 weeks (**Figure 4J**). EFS was further extended in mice that received IR treatment of 10 Gy total, with 5 out of 9 mice demonstrating complete response (CR) to the therapy (**Figure 4K**). However, it was the combination of oral trametinib treatment alongside IR therapy that culminated in all 10 mice displaying a prolonged CR (unpaired t-test compared to IR group CR mice, *p* = 0.0003), extending their EFS to 11.5 weeks (*p* < 0.0001, **Figure 4L**).

Finally, the second PDX, SJRHB000026_X1 control tumors had a slightly lower EFS (1.8 weeks) compared to the other FN-RMS xenografts when left untreated (**Figure 4M**). Similarly, the mice that received trametinib single-agent therapy responded poorly to single-agent treatment, displaying a shorter EFS compared to the other FN-RMS trametinib-treated groups, with most tumors reaching 4 times their initial volume during their trametinib treatment (**Figure 4N**). SJRHB000026_X1 PDXs that received IR therapy (20 Gy total) had EFS extended to an average of 7.9 weeks, with half of the mice showing CR where tumors were not palpable for an average of 1 week (**Figure 4O**). All but one of the SJRHB000026_X1 mice that received the combination therapy had a CR, nearly double the number of mice in the IR alone group (unpaired t-test compared to IR group CR mice, *p* = 0.0186). Additionally, the average EFS in the combination group was extended to 10.9 weeks, demonstrating a significant delay in relapse compared to IR treatment alone (*p* < 0.0001, **Figure 4P**). Together, these data suggest that the combination of a MEK inhibitor with IR therapy is more beneficial to slowing tumor growth and extending EFS than IR alone in FN-RMS and corresponds with previous SNAI2-knockdown/IR xenograft studies, implicating that SNAI2 downstream of MEK signaling is both a biomarker and a determinant of resistance to IR in RAS-mutant FN-RMS tumors.

## Discussion

In FN-RMS, there is a high incidence of RAS mutations, making RAS signaling an attractive therapeutic target for this childhood cancer. Mutations that activate RAS signaling are detected in all three RAS isoforms in FN-RMS, with NRAS mutations at codon Q61 being most frequent^1, 3^. Additionally, activating mutations in FGFR4 or PI3K, as well as loss of NF1, that activate RAS signaling are also observed^1–3^. Three recent studies that used either high dose MEK inhibitor trametinib, vertical inhibition of the RAS/RAF/MEK/ERK signaling pathway using RAF/MEK inhibitor combinations, or ablation of mutant NRAS suggest that RAS mutant FN-RMS tumors are addicted to mutant RAS signaling^13, 14, 27^. Additionally, they reveal a heterogeneity in response in FN-RMS when RAS signaling is lost, where tumor cells undergo myogenic differentiation, apoptosis, and a G1 cell cycle block. However, these reports and our current study also reveal that treatment of FN-RMS tumor xenografts with clinically relevant exposures of trametinib may not be sufficient to achieve robust responses. Moreover, studies targeting of MEK with single agents in RAS-dependent adult cancers have shown limited activity in patients^28^. In our current study, we sought to address if combining a clinically relevant trametinib dose with IR, a standard of care therapy for FN-RMS, would be an effective treatment. Additionally, we investigated whether SNAI2 downstream of RAS/MEK signaling can both prevent myogenic differentiation and protect FN-RMS tumors from radiation-induced apoptosis.

Our findings show that SNAI2 is a critical transcription factor downstream of RAS/MEK signaling that, in the context of radiation therapy, blocks myogenic differentiation and protects FN-RMS tumor cells from mitochondrial apoptosis. In contrast, we find that MEK signaling effects on the cell cycle are independent of SNAI2. Importantly, in preclinical models, we show that using a clinically relevant low dose of trametinib is sufficient to significantly reduce SNAI2 expression and when combined with IR results in complete responses in many of the tumor-burdened mice from the combination treatment group and a significant delay in EFS. This effect is particularly significant, as we find in our xenograft and PDX models, treatment with 1 mg/kg trametinib results in only ∼10% partial responses across all xenografts tested (3 of 30 mice); moreover, in one of the PDXs, tumors reached 4x their initial volume during their 28-day trametinib treatment. There are three important aspects to our findings related to the responses of FN-RMS tumors using the combination of MEK inhibitors and IR. First, there is an apparent heterogeneity in response to radiation after MEK inhibition. Second, MEK inhibitor, trametinib, significantly reduces SNAI2 expression, creating a clinically relevant way to reduce SNAI2 levels in tumors. Lastly, our findings have potential relevance in treating other RAS-mutant sarcomas.

When treated with MEK inhibitor trametinib and IR, FN-RMS cells undergo robust myogenic differentiation, apoptosis, and G1/S cell cycle block. It is important to note that in our experiments we do not use serum starvation, but rather maintain regular growth (10% FBS) conditions. Low serum assays are commonly used in the muscle and rhabdomyosarcoma field to determine effect of genes, pathways, or chemicals on induction of myogenic differentiation^21, 24–26^. Furthermore, when FN-RMS cells are treated with trametinib at low concentrations (5–10 nM), which is similar to concentrations tolerated in patients, the cells show some myogenic differentiation, but minimal apoptosis, even though our analyses show that these cells express elevated levels of BIM and are primed for apoptosis. These results are consistent with our previous studies, where we showed that a low dose of trametinib (5 nM) induces very little differentiation and almost no apoptosis *in vitro*. Additionally, SMS-CTR xenografts, known to be sensitive to MEK inhibition, only show partial responses and progressive disease post 1 mg/kg trametinib treatment *in vivo*^13^. However, robust myogenic differentiation and apoptosis were observed in FN-RMS cells and tumors *in vitro* and *in vivo* only after treatment with high trametinib concentrations, combining low-dose MEK inhibition with RAF inhibition, or ablating mutant NRAS with CRISPR/Cas9 tools^13, 14, 27^. Additionally, our data show that the response to MEK inhibition can vary between RAS-mutant PDX models, where some PDXs and xenografts have partial to poor responses to treatment with 1 mg/kg trametinib, which has been seen in other studies as well^14^. Similarly, when treated only with IR, while some apoptosis is observed, very little myogenic differentiation is induced in FN-RMS *in vitro* assays. However, the combination of MEK inhibitor and IR results in the robust initiation of myogenic differentiation and apoptosis. This not only results in the loss of colony formation post IR but also complete tumor responses in the majority of combination-treated mice and a consequent delay in event free survival by 2–4 weeks. Thus, myogenic differentiation and apoptosis are part of the spectrum of responses to MEK inhibitor and IR treatment. Our data suggest that combining MEK inhibitor treatment with standard of care IR treatment results in similar beneficial effects as combining trametinib with IGF1 inhibitor or combining trametinib with RAF inhibitors, even in tumors that have partial or poor responses to either clinically relevant MEK inhibitor concentrations or radiation alone^14^. Oncogenic RAS/MEK signaling has been shown to support proliferation, self-renewal, and prevent differentiation and apoptosis in RAS-mutant FN-RMS *in vitro* and *in vivo*. However, an understanding of how or if RAS/MEK signaling coordinates downstream effectors to determine whether tumor cells undergo self-renewal, myogenic differentiation, apoptosis, or exit the cell cycle remains fragmentary. This is of particular importance, as effector genes/proteins are likely also to be biomarkers of response to therapy in patients with RAS-mutant FN-RMS. Elegant genomic studies coupled with drug screens and experiments *in vitro* and *in vivo* by Yohe *et al*. provide insights on epigenetic regulation by downstream RAS effector, ERK2, on preventing myogenic differentiation^14^. ChIP-seq analyses of ERK2, in combination with analyses of other histone marks in FN-RMS SMS-CTR cells, showed that ERK2 is often bound close to the promoter and or at transcriptional start sites of genes regulating myogenesis and through interactions with RNA pol II causes transcriptional stalling and loss of full-length gene transcription. Furthermore, in FN-RMS cells, ERK2 is often deposited at myogenic transcription factors and structural proteins required for terminal differentiation, which results in repressed expression of these genes, causing cancer cells to resemble fetal skeletal muscle. Consequently, when these cells are treated with MEK inhibitor, ERK2 is lost, leading to induction of *MYOG, MYOD1, MEF2A, MEF2C,* and other myogenic structural genes and thereby releasing the differentiation block. Additionally, MEK inhibition results in increased chromatin accessibility in genes involved in terminal myogenic differentiation. ChIP-seq analyses for critical muscle differentiation transcription factors MYOG and MYOD in control versus trametinib-treated SMS-CTR cells found that MYOG and MYOD binding at enhancers and super enhancers results in the activation of a myogenic differentiation program^14, 19^.

Other studies have identified several RAS-modulated genes and proteins including CDKN1A, DNA-PKCs, Nanog, and stem cell genes that are either lost or upregulated when RD cells are treated with MEK inhibitor U0126^29–33^. However, how these genes are regulated downstream of RAS signaling or whether they contribute to resistance in RAS mutant FN-RMS remains to be determined. Our studies provide insights into how the downstream RAS/MEK-regulated effector SNAI2 can prevent both myogenic differentiation and apoptosis in response to radiation. Our previous and current experiments show that RAS/MEK signaling is required to stabilize SNAI2 protein rather than being required for its transcription^19^. Moreover, the differentiation program associated with ablation of SNAI2 in FN-RMS RD and SMS-CTR cells significantly overlaps with the program repressed by RAS/MEK signaling. We have additionally shown that the loss of SNAI2 in trametinib-treated cells results in the loss of SNAI2 binding to gene enhancers and super enhances required for myogenic differentiation^19^. Independently, we have demonstrated that SNAI2, through direct repression of pro-apoptotic BIM expression, prevents apoptosis post treatment with IR in both FN- and FP-RMS tumors. Thus, SNAI2 repression of BIM is likely shared across other sarcomas and tumors that express high SNAI2. RAS/MEK signaling is also known to turn over BIM through phosphorylation and subsequent degradation via the 16s proteasome complex^20^. Using classic rescue experiments, we show that expressing SNAI2 in cells where RAS/MEK signaling is inhibited can both prevent myogenic differentiation, as well as prevent BIM-mediated apoptosis after radiation exposure. Importantly, our data clearly show that a finite treatment combining MEK inhibition and IR is sufficient to result in complete tumor responses and significantly extended EFS, long past the conclusion of therapeutic intervention. Thus, SNAI2 is a major effector of RAS-MEK effects on preventing IR-induced myogenic differentiation and apoptosis, while there are likely other effectors modulating cell cycle. In our experiments, we show that loss of SNAI2 also significantly reduces FN-RMS RD and SMS-CTR sphere formation *in vitro*^19^, suggesting additional roles for SNAI2 in expanding self-renewal by yet to be determined mechanisms. Moreover, a recent study showed that Snai2 is required to prevent aging in skeletal muscle, and the loss of Snai2 in mice results in the loss of repopulating muscle stem cells due to increased number of cells undergoing senescence^34^.

Our studies showing a RAS/MEK/SNAI2 program preventing myogenic differentiation and a RAS/MEK/SNAI2/BIM program preventing apoptosis might also be relevant to other sarcomas and RAS-mutant tumors. For example, recent reports in RAS-mutant pancreatic cancer and melanoma showed that resistance to MEK inhibitor treatment can be acquired due to the upregulation of SNAI2^35, 36^. SNAI2 high cells re-establish proliferation and become more metastatic, while ablating SNAI2 re-sensitizes these cells to MEK inhibitor. Trametinib and other MEK inhibitors are currently being tested in combination treatments with IR in other RAS- mutant tumors^15, 16, 37^. Furthermore, analyses of The Cancer Genome Atlas (TCGA) data sets show that sarcomas generally express high levels of SNAI2, and IR is commonly used as standard of care therapy for sarcomas. For example, osteosarcomas express some of the highest levels of SNAI2, and IR is not commonly used to treat this disease; in contrast, Ewing sarcoma, another tumor found in the bone, express low SNAI2 and are known to be extremely sensitive to IR^38^. However, whether SNAI2 is the downstream effector or could be used as a biomarker for response remains to be explored. In conclusion, we show that inhibition of the RAS/MEK/SNAI2 axis with a low dose of trametinib in combination with IR is sufficient to ablate SNAI2 expression and consequently result in robust myogenic differentiation and apoptosis (**Figure 5**), leading to delayed tumor recurrence.

**Figure 5.**
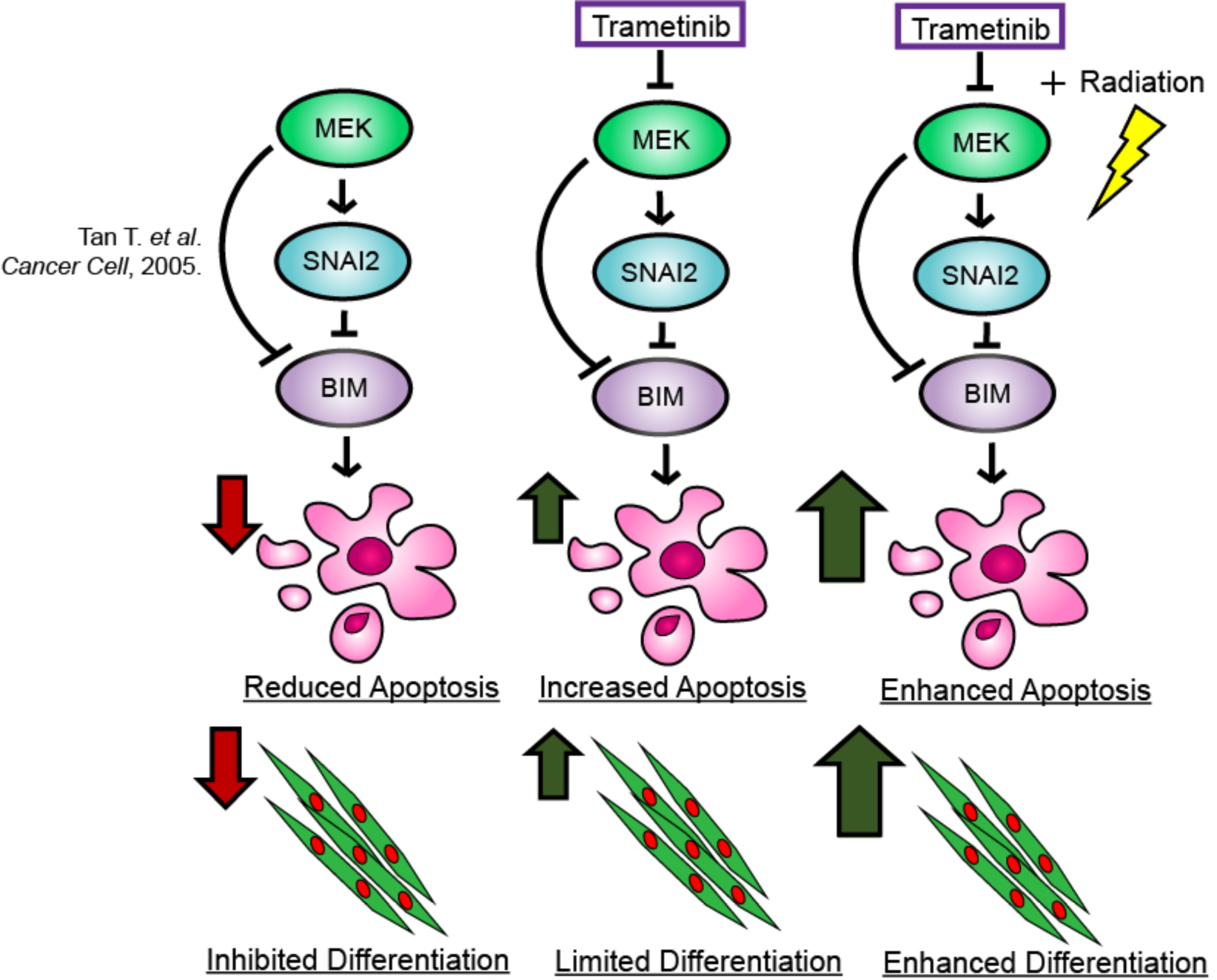
Proposed mechanism of inhibiting the MEK/SNAI2 pathway to sensitize FN-RMS to IR-induced apoptosis and myogenic differentiation.

## Figure Captions

**Supplemental Figure 1.**
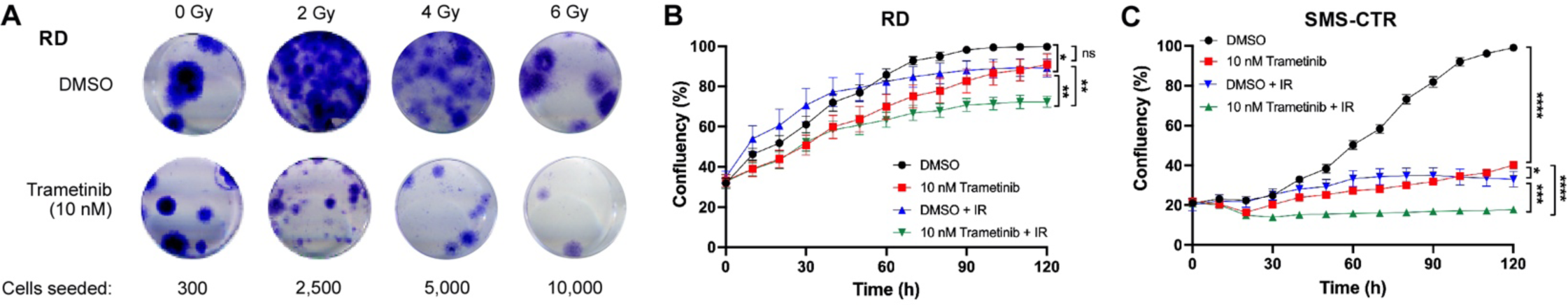
A. Representative colony formation wells from RD cells treated with DMSO or 10 nM trametinib then exposed to the indicated IR dose. B, C. Average percent confluency of RD (A) and SMS-CTR (B) cells after DMSO and trametinib treatment, with and without IR exposure. Error bars represent ± 1 SD. \**p* < 0.05, \*\**p* < 0.01, \*\*\**p* < 0.001, \*\*\*\**p* < 0.0001 by one-way ANOVA with Dunnett’s multiple comparisons test.

**Supplemental Figure 2.**
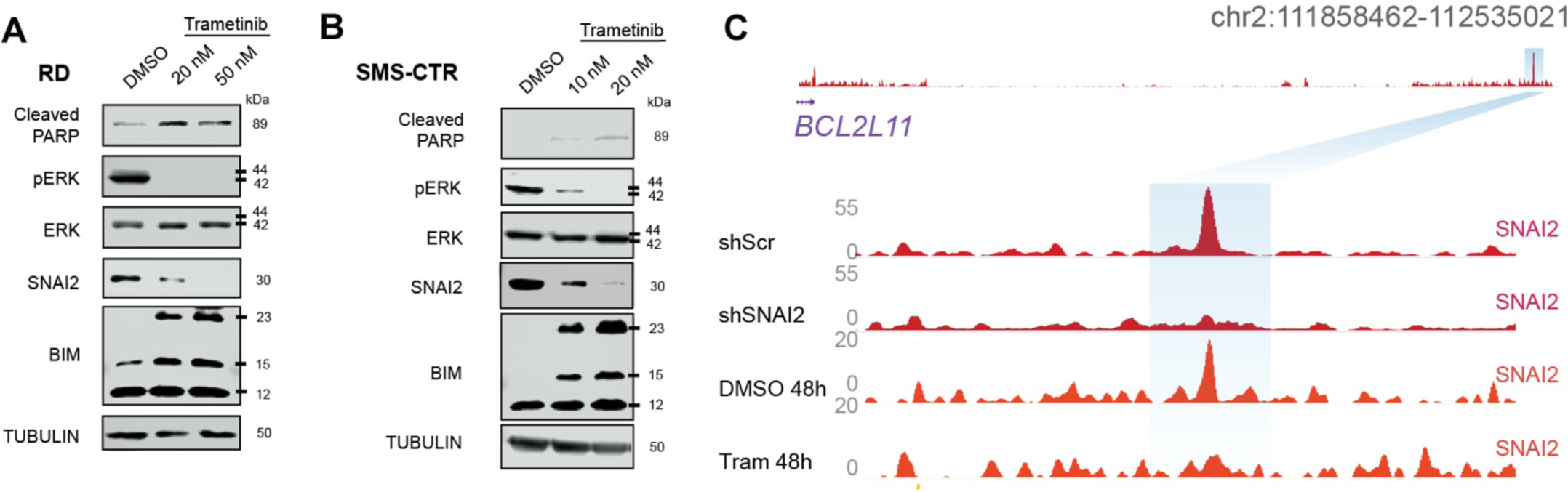
A, B. Cleaved PARP, pERK, total ERK, SNAI2, and BIM protein expression immunoblots in DMSO- and trametinib-treated RD (A) and SMS-CTR (B) cells treated with DMSO and increasing doses of trametinib for 24h. C. ChIP-seq SNAI2 binding peaks of the downstream enhancer region of *BCL2L11* in SMS-CTR cells treated with DMSO or trametinib (10 nM) for 48h, compared to SNAI2 binding peaks seen in RD shScr or shSNAI2 cells.

**Supplemental Figure 3.**
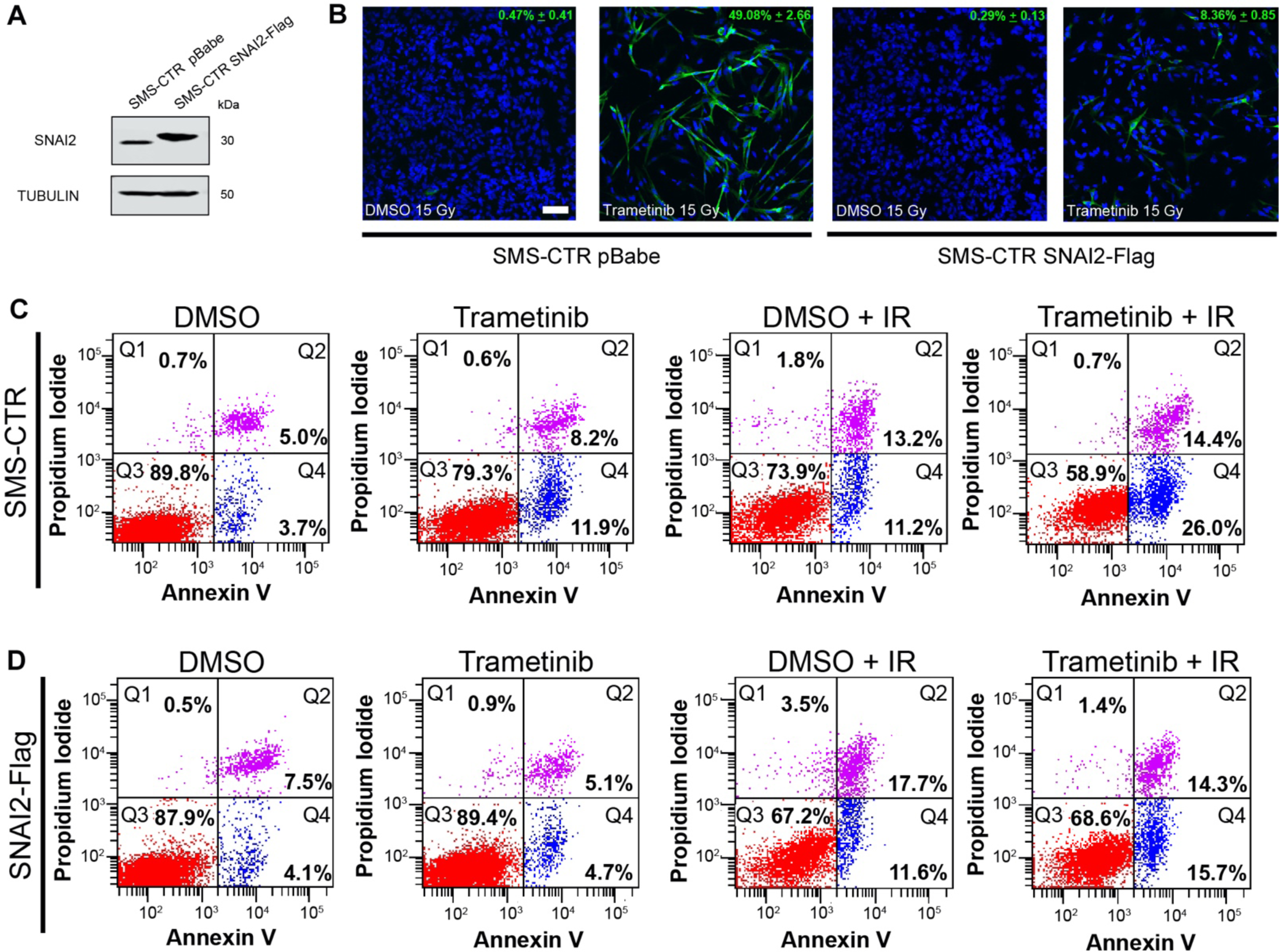
A. SNAI2 protein expression in SMS-CTR pBabe and SNAI2-Flag overexpressing cell lines. B. Representative confocal microscopy images of SMS-CTR pBabe and SNAI2-Flag cells treated with DMSO or trametinib after IR exposure, immunostained with myogenic differentiated myosin MF20 antibody. Scale bar = 100 μm. C. Flow cytometry plots showing propidium iodide vs. annexin V staining of DMSO- and trametinib-treated SMS-CTR pBabe cells after non-IR and IR conditions. D. Flow cytometry plots showing propidium iodide vs. annexin V staining of DMSO- and trametinib-treated SMS-CTR SNAI2-Flag cells after non-IR and IR conditions.

**Supplemental Figure 4.**
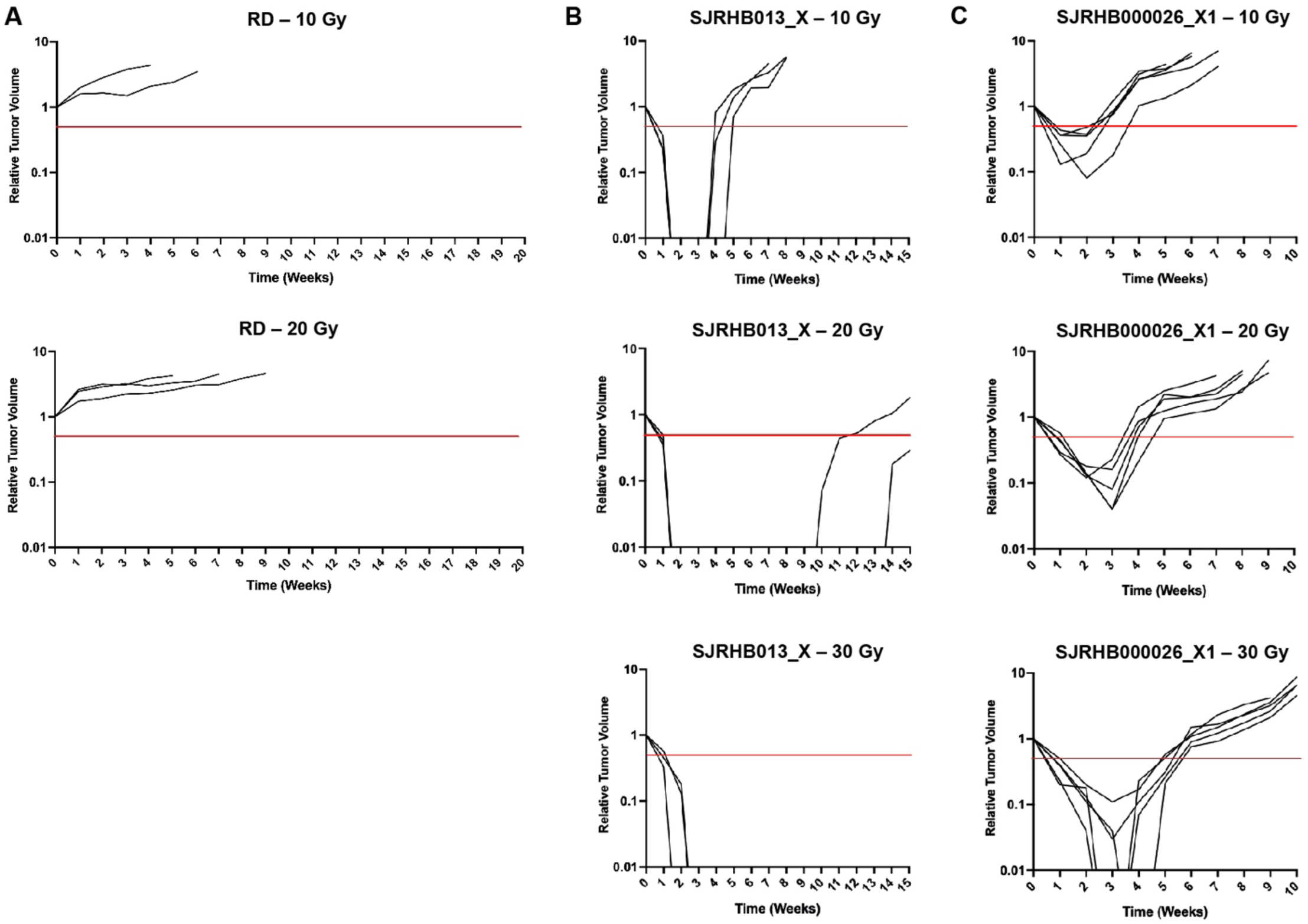
A–C. Relative tumor volume (RTV) of RD (A), SJRHB013_X (B), and SJRHB000026_X1 (C) tumors receiving increasing doses of IR (10–30 Gy) for IR sensitivity studies. Red line depicts 0.5 RTV.

**Supplemental Table 1.**

| **FN-RMS Cell Line** | **RAS Mutation** | **TP53 Status** |
| --- | --- | --- |
| RD | NRAS<sup>Q61H</sup> <sup>39</sup> | R248W mutation<sup>40</sup> |
| SMS-CTR | HRAS<sup>Q61K</sup> <sup>39</sup> | ΔNt 1236-1239<sup>40,41</sup> |
| Rh36 | HRAS<sup>Q61K</sup> <sup>42</sup> | Wild type<sup>41</sup> |
| **FN-RMS PDX** |  |  |
| SJRHB013 X | NRAS<sup>Q61K</sup> <sup>38</sup> | Wild type<sup>38</sup> |
| SJRHB000026 X1 | HRAS<sup>G13R</sup> <sup>38</sup> | Wild type<sup>38</sup> |

**Supplemental Table 2.**

| **REAGENT/RESOURCE** | **SOURCE** | **IDENTIFIER** |
| --- | --- | --- |
| phospho ERK | CST | Cat# 4370, RRID:AB_2315112 |
| ERK | CST | Cat# 9102, RRID:AB_330744 |
| Slug (C19G7) | CST | Cat# 9585, RRID:AB_2239535 |
| SNAI1 (L70G2) | CST | Cat# 3895s, RRID:AB_2191759 |
| BIM (C34C5) | CST | Cat# 2933, RRID:AB_1030947 |
| Cleaved PARP (Asp214) (D64E10) | CST | Cat# 5625, RRID:AB_10699459 |
| GAPDH | CST | Cat# 2118, RRID:AB_561053 |
| β-Tubulin | Abcam | Cat# Ab6046, RRID:AB_2210370 |
| HRP anti-rabbit | CST | Cat# 7074, RRID:AB_2099233 |
| HRP anti-mouse | GE Healthcare | Cat# NA93IV, RRID:AB_772210 |
| Myosin Heavy Chain (MF20) | DSHB | Cat# MF 20, RRID:AB_2147781 |
| Alexa Flour 488 goat anti-mouse | Invitrogen | Cat# A11029, RRID:AB_138404 |

